# Diffusion-enhanced spatio-temporal models for estimating spatial expansion

**DOI:** 10.64898/2026.09.21.749344

**Authors:** Hsing-Han Wu, James T. Thorson, Yi Chang

**Affiliations:** Department of Oceanography, National Sun Yat-sen University, Kaohsiung, Taiwan; DTU Aqua, National Institute of Aquatic Resources, Technical University of Denmark, Kongens Lyngby, Denmark; Resource Ecology and Fisheries Management Division, Alaska Fisheries Science Center, National Marine Fisheries Service, Seattle, Washington, USA; Graduate Institute of Marine Affairs, National Sun Yat-sen University, Kaohsiung, Taiwan

## Abstract

Spatial expansion is a common feature of many ecological processes, including biological invasions, range shifts, population recovery, and the spread of pathogens and human activities. Although such expansion often arises from diffusive processes, there remain few general statistical methods to quantify these dynamics. Here, we present a diffusion-enhanced spatio-temporal model (DESTM) that approximates a diffusive process using Gaussian Markov random fields (GMRF). The model works by incorporating the statistical interaction between spatial diffusion and temporal lags into a joint precision structure of the latent field. To demonstrate, we simulate data from a diffusion movement process using a continuous-time Markov chain (CTMC), and then approximate this mechanistic process using the DESTM. We show that separable spatio-temporal models can lead to large bias in estimates of the local density. In contrast, the DESTM did not appear to bias inferences, and reverted to the separable model when diffusion is not apparent. Finally, we apply the model to two large-scale examples of spatial expansion: (1) the historical spread of the Japanese distant-water longline fleet across the Pacific, representing anthropogenic spatial expansion in the ocean, and (2) the recovery and geographic expansion of Bald Eagle populations across North America. Estimates from both case studies, supported by model comparisons and ecological plausibility, suggest that the DESTM provides a better description of the underlying expansion dynamics, although improvements in forecast performance were more apparent for the Japanese longline fleet. We highlight that the model provides a fast and efficient framework for modeling spatial expansion dynamics, with the potential to represent increasingly complex ecological dynamics through future extensions. We propose that future research explore this framework as a quantitative tool to explain how spatial expansion occurs.

## 1. Introduction

Spatial expansion is a fundamental process in ecology, underlying biological invasions, climate-driven range shifts, population recovery, and the spread of pathogens and human activities. Across spreading populations, expansion is generally shaped by the interplay between two key processes: dispersal and demography (Chuang and Peterson, 2016; Miller et al., 2020). Dispersal involves departure from occupied areas, movement across the landscape, and settlement in unoccupied locations (Chuang and Peterson, 2016; Clobert et al., 2009). Local demographic processes, in turn, determine whether individuals survive and reproduce following arrival and whether newly colonized populations become established and persist under local environmental and biotic conditions (Hardie and Hutchings, 2010; Osland et al., 2021; Pigot et al., 2010). The combined effects of these processes can cause population distributions to expand across space over time (Kubisch et al., 2014; Miller et al., 2020).

In ecological modelling, species distribution models (SDMs) are widely used to understand the relationship between species and their environment, and to predict distributions across space and time (Elith and Leathwick, 2009). For spatial expansion specifically, static, correlative SDMs may face limitations arising from their underlying assumptions. These static SDMs relate species presence to abiotic and biotic factors to estimate habitat suitability, often assuming that species tend to occur in environmentally favorable areas without explicitly accounting for dispersal processes or limitations (Guisan and Thuiller, 2005; Klaassen et al., 2026). However, the presence or absence of an expanding population in an area does not necessarily reflect environmental suitability, because its distribution may be limited by dispersal limitation or geographic barriers (Gaylord and Gaines, 2000; Pecl et al., 2017). Ignoring these dynamics may therefore limit our understanding of how population distributions change (Pinsky et al., 2020).

Modelling spatial expansion therefore requires moving beyond static SDMs to account for changes in species distributions through time, e.g., mechanistic models and spatio-temporal models (STMs). Mechanistic models explicitly incorporate biological processes, such as survival, reproduction, and movement, to predict species distributions (Kearney and Porter, 2009; Kearney et al., 2010). To model spatial expansion dynamics, mechanistic models can specify diffusive process that translate individual movement into population-level changes in density through flows among neighboring locations over time (Hefley et al., 2017b). Specifying this process, however, requires prior knowledge or assumptions about movement and its environmental drivers, which may not always be available or well understood (Dormann et al., 2012; Hefley et al., 2017a). Alternatively, spatio-temporal models (STMs) can be used to estimate changes in species distributions directly from observations across space and time (Thorson and Kristensen, 2024). These models instead use latent spatial or spatio-temporal random fields to account for dependence across locations and times and variation not explained by environmental covariates (Anderson et al., 2025; Thorson, 2019). In many common STMs, temporal dependence carries the latent spatial field forward at the same locations, while spatial dependence is modelled separately within each time period (often called separable STMs). Although STMs can describe changes in species distributions through time, the estimated latent field is not necessarily linked to the ecological processes driving spatial expansion (Cressie and Wikle, 2011; Hefley et al., 2017a).

Latent fields in STMs can, however, be constructed to represent particular processes. Recent developments in spatial statistics represent diffusion within the latent fields of STMs by deriving the precision matrices of GMRFs (hereafter, diffusion-enhanced GMRFs) from diffusion-based process equations. For example, Lindgren et al. (2024) extended Gaussian Matérn fields using the stochastic partial differential equation (SPDE) approach to couple temporal change with spatial diffusion.

Alternatively, Thorson (2026) used a graphical Gaussian model to describe spatial diffusion and temporal dependence among latent states and map these relationships into the GMRF precision matrix. Diffusion-enhanced GMRFs retain the computational advantages of sparse GMRFs while defining latent dependence in terms of diffusion through space and time. However, the potential of diffusion-enhanced GMRFs for modelling expanding distributions remains unexplored in ecology.

In this study, we introduce a diffusion-enhanced spatio-temporal model (DESTM) that approximates spatial diffusion processes within a Gaussian Markov random field framework. The approach links spatial diffusion and temporal dependence through a joint precision structure, allowing neighboring locations at previous times to influence the current spatial distribution (a non-separable STM, with the separable form included as a special case). To evaluate the DESTM, we conduct simulation experiments using a continuous-time Markov chain (CTMC) movement process to examine model performance in predicting CTMC-generated diffusive dynamics. Finally, we apply the approach to two case studies: (1) the historical expansion of the Japanese distant-water longline fleet across the Pacific Ocean and (2) the recovery and geographic expansion of Bald Eagle populations across North America following widespread population declines. We assess model performance in both case studies using predictive skill and ecological plausibility.

## 2. Methods

### 2.1. Diffusive process

Here, we develop a statistical approximation of spatial expansion that remains computationally tractable for large spatio-temporal datasets. We build on the path-matrix approach of Thorson (2026), which combines spatial diffusion, temporal dependence, and their interaction to describe diffusive dynamics. We adapt this structure to estimate diffusive dynamics and predict spatial expansion from observed data. In the following, we define the spatial and temporal operators underlying the model and then show how the resulting process can be written as a Gaussian Markov random field (GMRF).

We define neighboring relationships among the *S* locations in a two-dimensional domain using an *S × S* adjacency matrix **A**:

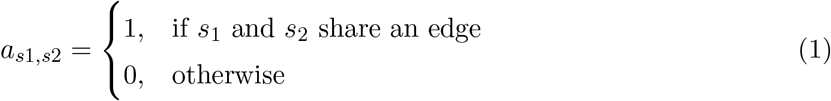

We then row-standardize this adjacency matrix by dividing each row by its number of neighboring locations, such that the weights of neighboring locations sum to one:

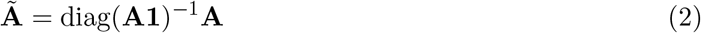

where **1** is an *S*-length vector of ones, such that **A1** gives the number of neighboring locations for each location. To represent local diffusion among neighboring locations, we transform row-standardized adjacency matrix *Ã* into a spatial diffusion operator:

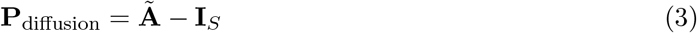

where **I**_*S*_ is an *S × S* identity matrix. This diffusion operator **P**_diffusion_ represents the difference between the density of neighboring locations and the density at a focal location, allowing density to spread from relatively high-density locations toward adjacent lower-density areas. Because each row of **P**_diffusion_ sums to zero, local losses are balanced by gains among neighboring locations.

To represent spatial expansion through time, we next extend diffusion within a single time period by incorporating temporal dependence, allowing state at one time period to influence that at the subsequent period. Specifically, we define a first-order temporal lag operator **P**_lag1_, where non-zero elements link each time period *t* to the previous period *t −* 1:

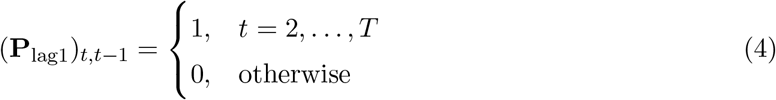

We then combine the spatial diffusion and temporal lag operators using Kronecker product to define a joint spatio-temporal path matrix **P**_joint_ for spatio-temporal diffusion processes:

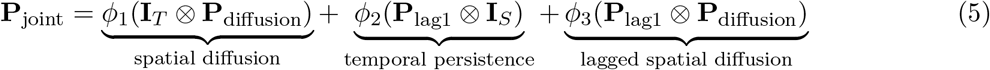

where **I**_*T*_ is an *T × T* identity matrix, *⊗* is Kronecker product, and *ϕ*_1_, *ϕ*_2_ and *ϕ*_3_ are coefficients controlling the relative strength of spatial diffusion, temporal lag, and lagged spatial diffusion, respectively. For the non-separable DESTM, *ϕ*_3_ is estimated freely in Equation 5. This expression can also revert to a separable spatio-temporal structure, when *ϕ*_3_ = *−ϕ*_1_*ϕ*_2_, in which spatial diffusion and temporal dependence operate independently through time (hereafter, separable DESTM; see Example 2 in Thorson, 2026). In the separable form, the expected latent state *x*_*s,t*_ depends on the previous state at the same location *x*_*s,t−*1_, whereas, in the non-separable form, it also depends on previous states 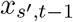 at locations *s*^*′*^ neighboring *s* through the spatial operator.

After defining the joint spatio-temporal operator **P**_joint_, we use it to specify the spatio-temporal dependence underlying the latent diffusion process. This process can be expressed as a simultaneous equation:

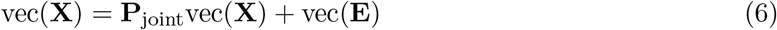

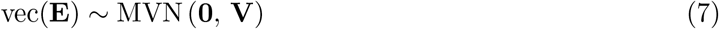

where **X** is a *S×T* matrix of latent states containing *x*_*st*_ such that vec(**X**) stacks columns into a state vector with length *ST*, **E** is the *S × T* matrix of spatio-temporal process errors, and **V** is a diagonal variance matrix controlling the magnitude of unexplained variation. This latent process is described as the combination of structured spatio-temporal dependence implied by diffusion dynamics and residual variation not captured by the expected process. We can rewrite this relationship as:

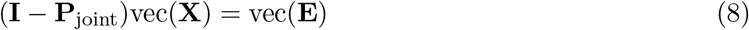

where (**I** *−* **P**_joint_)vec(**X**) represents departures from the expected spatio-temporal structure implied by diffusion and temporal lag. In this form, the operator (**I** *−* **P**_joint_) measures the discrepancy between the latent state and the expected state predicted from neighboring locations and previous time periods. Because unexplained variation follows a multivariate normal distribution, weighted departures from the expected spatio-temporal process can be represented through the precision matrix:

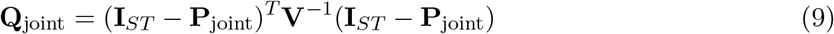

**I**_*ST*_ is an *ST ×ST* identity matrix and the quadratic form (**I**_*ST*_ *−***P**_joint_)^*T*^ **V**^*−*1^(**I**_*ST*_ *−***P**_joint_) represents weighted departures from the expected spatio-temporal process, with **V**^*−*1^ controlling the strength of penalization. Larger values of **V** allow greater departures from the expected diffusion dynamics, whereas smaller values impose stronger adherence to the spatio-temporal structure defined by **P**_joint_. The latent state vector therefore follows a Gaussian Markov random field (GMRF; Rue and Held, 2005):

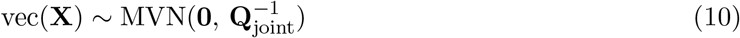

Because **P**_joint_ is sparse and **V** is diagonal, the resulting precision matrix **Q**_joint_ is also sparse, allowing computationally tractable estimation for large spatio-temporal datasets.

### 2.2. Simulation experiment

We conduct two simulation experiments to examine the following three questions:

1. can the DESTM recover and forecast spatial expansion generated by a mechanistic diffusion process?
2. how robust is model performance to different expansion scenarios?
3. how do separable and non-separable DESTMs perform when fitted to different spatio-temporal processes?

To answer the first two questions, we use a continuous-time Markov chain (CTMC) model to simulate diffusion dynamics under different scenarios, including point- and strip-source initial conditions, varying diffusion rates, and observation coverage. We then fit both separable and non-separable models to the simulated data and evaluate how well the models can recover and forecast the simulated diffusion process. To answer the third question, we conduct a self and cross simulation experiment in which data are generated from both separable and non-separable spatio-temporal processes and fit using both models.

#### 2.2.1 CTMC diffusion experiment

We first test the DESTM using data generated from a CTMC diffusion simulation model. Specifically, we evaluate model performance across multiple expansion scenarios, including point- and strip-source initial conditions, three levels of observation coverage (30%, 70%, and 100%), and varying diffusion rates (*δ* = 0.3, 0.5, and 0.8). The CTMC simulation model involves generating log-density *x*_*s,t*_ at location *s* (*s ∈ {*1, …, *S}*) and time *t* (*t ∈ {*1, …, *T}*) over a square spatial domain discretized into a 20 *×* 20 grid (*S* = 400) across *T* = 10 time steps.

To represent local movement among neighboring grid cells, we use the adjacency matrix A defined in Equation 1 to construct a diffusion matrix **D** = *δ*[**A***−*diag(**A1**)] representing instantaneous movement rates. The adjacency matrix **A** is scaled by *δ* to give a movement rate of *δ* between neighboring cells, while each diagonal element is the negative total rate of movement out of the corresponding cell. To propagate the log-density field through time, we integrate these movement rates over one time interval using the matrix exponential:

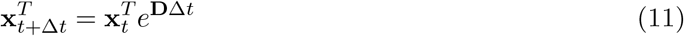

We consider two initial scenarios to represent different forms of ecological spread. In the point-source scenario, expansion originates from a localized introduction point positioned at the center of the spatial domain, representing localized expansion from a single introduction source. In the strip-source scenario, expansion is initialized along one boundary of the domain to represent directional expansion entering from the edge of the system. For each initial scenario, diffusion is initialized at t = 1 using a total initial intensity of 100 distributed according to the specified source configuration, after which dynamics are projected forward through time (Fig. 1).

**Figure 1.**
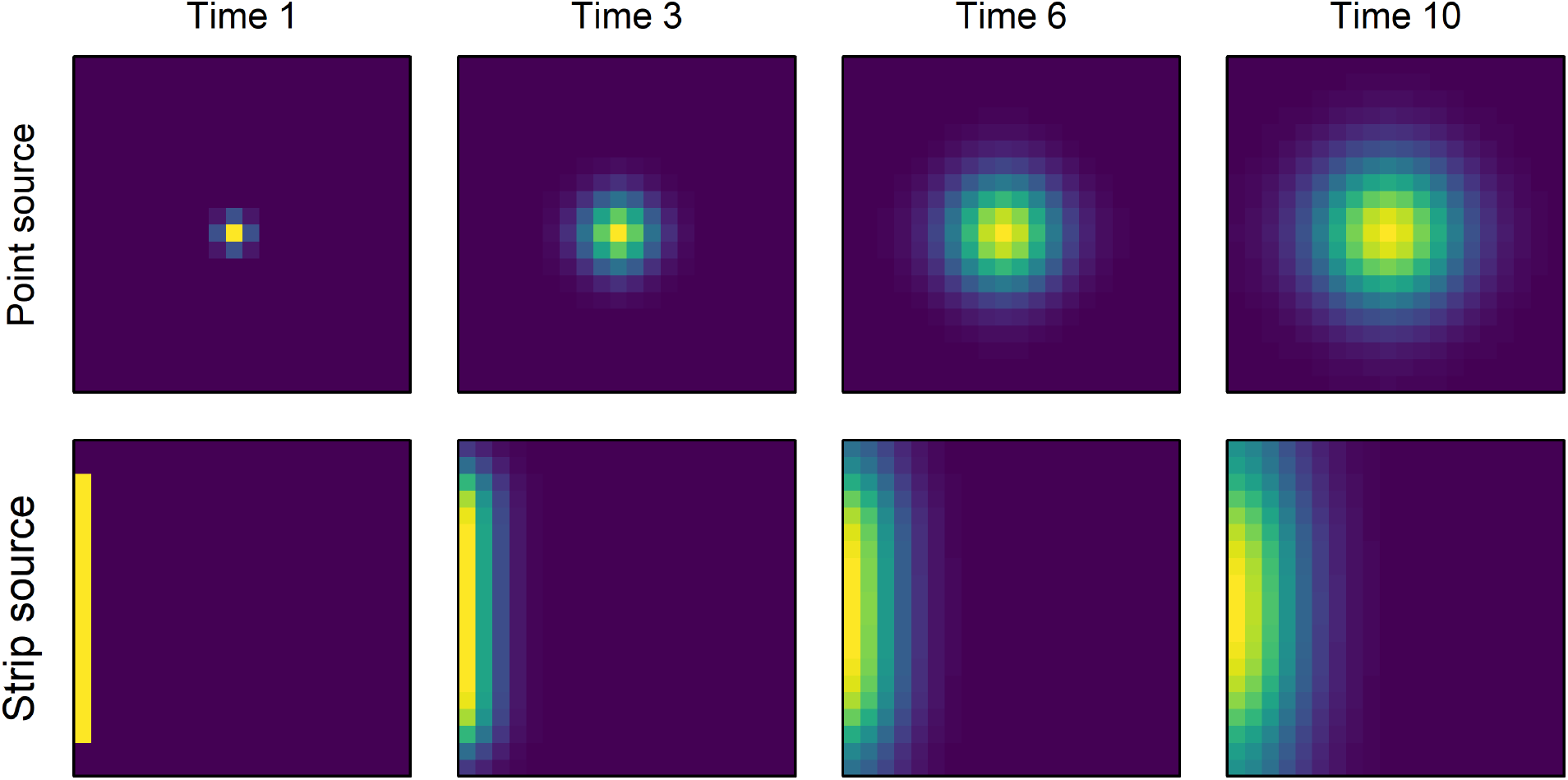
Visualizing CTMC diffusion dynamics under point-source (top row) and strip-source (bottom row) initial conditions at *t* = 1, 3, 6, and 10. Simulations are illustrated using an intermediate diffusion coefficient (*δ* = 0.5). Color scales were mapped to the value range within each time step to visualize diffusion through time.

In addition to simulations assuming the full spatial distribution is perfectly observed, we also consider a sampling process to better resemble ecological datasets obtained through sampling or collection. Specifically, the simulated log-density field is first exponentiated to obtain relative intensities, which were then normalized to sampling probabilities, with samples at each time step generated from a multinomial distribution:

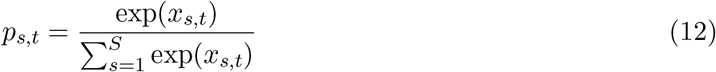

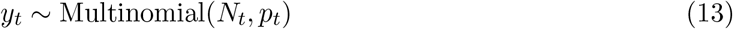

where *p*_*s,t*_ is the sampling probability at location *s* and time *t, p*_*t*_ is the vector of sampling probabilities across all grid cells, *N*_*t*_ denotes the sample size at time *t*, and *y*_*t*_ is the vector of sampled data across grid cells at time *t*. To mimic incomplete observation commonly encountered in ecological data, we evaluate three levels of observation coverage (30%, 70%, and 100%) under both fully observed and multinomial observation processes, where observations are randomly sampled at each time step. For each initial scenario (point- or strip-source) and diffusion rate (*δ* = 0.3, 0.5, and 0.8), simulations are conducted under all combinations of observation process and observation coverage. A more detailed illustration of the simulation scenarios, including the latent diffusion process, full samples, and samples with 70% and 30% observation coverage, is provided in Fig. S1. For each of 30 simulation replicates, we use the diffusion precision matrix **Q**_diffusion_ to specify separable and non-separable DESTMs using GMRFs, fit both models to simulated samples from *t* = 1–7, and project forward to *t* = 8–10 for out-of-sample forecasting. Model performance is then evaluated by comparing model predictions against the latent simulated truth at each time step using mean absolute error (MAE) across all grid cells.

#### 2.2.2 Self and cross simulation experiment

For question 3, we generate data from separable and non-separable DESTM operating models and fit both estimation models to evaluate model performance. We simulate 100 datasets for each combination of three diffusion scenarios (low, moderate, and high) and two operating models (separable and non-separable) using the same spatial domain and temporal extent described in the CTMC diffusion experiment. For every dataset, a new GMRF is simulated to generate realizations of the latent spatio-temporal random effects under an intercept-only model, without additional fixed effects. The spatial diffusion *ϕ*_1_, temporal lag *ϕ*_2_, and lagged spatial diffusion (interaction; *ϕ*_3_) are based on estimates from the fully observed point-source CTMC diffusion scenarios, with small random variation added across datasets within each diffusion scenario. Datasets are then simulated conditionally using a low-variance compound Poisson–Gamma Tweedie distribution. For each dataset, we fit both separable and non-separable estimation models using samples from *t* = 1–7 and forecast *t* = 8–10. We evaluate prediction and forecast performance by comparing model predictions with the simulated datasets using MAE.

### 2.3 Case study

To illustrate the application of the DESTM across contrasting forms of spatial expansion, we consider two case studies: (1) the historical spread of the Japanese distant-water longline fleet across the Pacific Ocean and (2) the recovery and geographic expansion of Bald Eagle populations across North America.

#### 2.3.1 Japanese distant-water longline fleet

To illustrate the application of diffusive processes in practice, we seek to approximate the spatial and temporal dynamics of a large-scale range expansion using the historical spread of the Japanese distant-water longline fleet across the Pacific as an example. We therefore compile publicly available longline catch and effort data from the Western and Central Pacific Fisheries Commission (WCPFC) and the Inter-American Tropical Tuna Commission (IATTC) for the period 1952–1974 (Inter-American Tropical Tuna Commission, 2026; Western and Central Pacific Fisheries Commission, 2026). We stop at 1974, given that later years include an increasing number of national fishing fleets, which are likely to have different dynamics than the Japanese longline fleet that is dominant during 1952 through the mid-1970s. We define a spatial domain, divided into 10° latitude by 10° longitude cells, and assume that all boundaries are reflective (i.e., no movement into or out of the domain). Fishing effort, measured as the total number of hook sets, is then summed for each cell in each year.

We fit separable and non-separable DESTMs to samples of fishing effort, *y*_*s,t*_, at location *s ∈ {*1, 2, …, *S}* for *S* = 139 spatial grids and discrete time interval *t ∈ {*1, 2, …, *T}* spanning *T* = 23 years. We assume that fishing effort follows a Tweedie distribution with a log link:

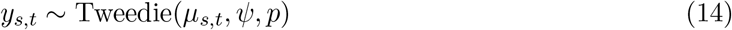

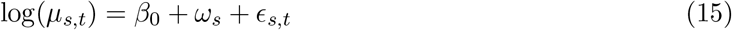

where *µ*_*s,t*_ is the expected fishing effort at location *s* and time *t*, and *ψ* and *p* are dispersion parameters controlling the magnitude and mean-variance relationship of the observation distribution. Here, *β*_0_ is intercept, *ω*_*s*_ represents the spatial effect at location *s*, and *ϵ*_*s,t*_ denotes the diffusion effect at location *s* and time *t*. We specify Gaussian Markov random fields (GMRFs) for both the spatial and diffusion effects:

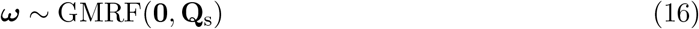

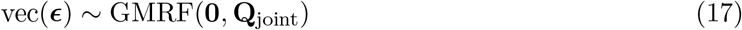

where **Q**_joint_ is the spatio-temporal diffusion precision matrix defined in Equation 9. For the spatial effect, we define the spatial precision matrix as:

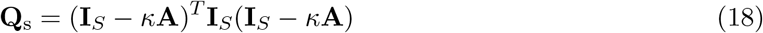

where **A** is the binary adjacency matrix defined in Equation 1, **I**_*S*_ is an *S × S* identity matrix, and *κ* controls the degree of spatial smoothing among neighboring cells.

Finally, we fit both separable and non-separable DESTMs using all available data (1952–1974). We compare models using marginal Akaike’s Information Criterion (mAIC; Zheng et al., 2024), deviance explained, mean absolute error (MAE), root mean squared error (RMSE), correlation (COR), and continuous ranked probability score (CRPS) (Gneiting et al., 2005). Results from the selected model are then examined to determine whether it can explain range expansion in the longline fishery observed in the early 1970s. We also conduct a predictive evaluation of model suitability. Specifically, we fit both DESTMs using a leave-future-out approach with a rolling window. For each year from 1957 to 1971, models are fitted using data from 1952 through year t and then used to predict dynamics for the following three years (*t* + 1 to *t* + 3). We then compare predictions with the observed data for these future years. Forecast performance is evaluated using MAE, RMSE, COR, and CRPS.

#### 2.3.2 Bald Eagle population recovery

For the second case study, we examine the recovery and geographic expansion of Bald Eagle populations across North America. We compile publicly available count data from the North American Breeding Bird Survey (BBS) for the period 1966–2025, excluding 2020 because BBS data were not available for that year (*T* = 59 years) (Ziolkowski et al., 2026). The study area includes the United States and Canada, including Alaska but excluding Hawaii. The dataset contains 143,679 records from 5,642 survey routes. We divide the study area into 5° latitude by 5° longitude cells and retain the *S* = 104 cells containing at least one BBS route. For each cell and year, we sum the total number of Bald Eagles recorded across routes and calculate sampling effort as the number of unique routes surveyed. Cell-years without surveyed routes are treated as missing rather than as observed zeros.

We fit both separable and non-separable DESTMs to the aggregated Bald Eagle count *y*_*s,t*_, at location (*s ∈ {*1, 2, …, *S}*) and year (*t ∈ {*1, 2, …, *T}*). We assume that the observed count follows a Poisson distribution with a log link:

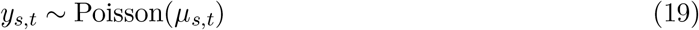

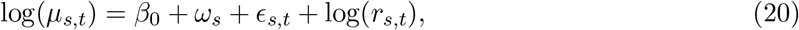

where *µ*_*s,t*_ is the expected Bald Eagle count and *r*_*s,t*_ is the number of routes surveyed within cell *s* during year *t*. log(*r*_*s,t*_) is included as an offset to account for variation in sampling effort among cells and years. As in the Japanese longline case study, *β*_0_ is the intercept, *ω*_*s*_ is the spatial effect, and *ϵ*_*s,t*_ is the diffusion effect. We use the same GMRF specifications for the spatial and diffusion effects described above.

We first fit the separable and non-separable DESTMs using all available observations from 1996 to 2025. We compare models using mAIC, deviance explained, MAE, RMSE, COR, and CRPS, following the same model-comparison procedure used for the Japanese longline case study. We also evaluate predictive performance using rolling-origin leave-future-out cross-validation with an expanding training period. For each iteration, models are fitted using data from 1966 through year *t* and then used to forecast years *t* + 1, *t* + 2, and *t* + 3. For example, the first fit uses data through *t* = 1985 to forecast 1986–1988, and the final fit includes data through 2016 and forecasts for 2017–2019. We do not extend the evaluation beyond 2019 because BBS data are unavailable for 2020. Forecasts are conditional on the observed number of BBS routes surveyed in each future cell-year. We evaluate the final fitted year and each of the three forecast horizons using MAE, RMSE, COR, and CRPS, with CRPS approximated using 2,000 samples from the Poisson predictive distribution. Metrics are calculated only for cell-years in which at least one route was surveyed.

To additionally evaluate predictions of whether Bald Eagles were observed, we classify surveyed cell-years with positive counts as observed presences and those with zero counts as observed absences. Under the Poisson observation model, the predicted probability of a positive count is calculated as 1 *−* exp(*−µ*_*s,t*_), conditional on the number of routes surveyed in each cell-year. We then compare these predicted probabilities between observed presences and absences for the final fitted year and each of the three forecast horizons, separately for the separable and non-separable models.

## 3. Results

### 3.1. Simulation experiments

We first visualized the diffusion dynamics generated by the mechanistic CTMC model andestimates from the separable and non-separable DESTMs to illustrate how the statistical model approximates the diffusion process (Fig. 2). In this example, the CTMC simulation was fully observed, with no process error or missing observations at an intermediate diffusion rate (*δ* = 0.5; Fig. 2, top row). Diffusion was generated from point-source (left panel) and strip-source (right panel) initial conditions. Under both source scenarios, the non-separable DESTM closely reproduced the expanding diffusion dynamics from the initial condition (*t* = 1; Fig. 2, left column), through an intermediate fitted state (*t* = 5; Fig. 2, middle column), to the three-step-ahead forecast (*t* = 10; Fig. 2, right column), whereas the separable model underestimated expansion during forecasting. We next evaluated model performance across all simulation scenarios to determine whether the visual patterns observed in Fig. 2 were consistent across simulation settings (Fig. 3). The results showed that the non-separable model provided lower estimation and forecast errors than the separable model, especially under incomplete observation coverage (30% and 70%; first and second columns), under process error (second and fourth rows), and during forecasting (grey-shaded panels). Across all three diffusion rates, estimation and forecast errors were generally lower for the non-separable model, although the differences between models did not consistently increase or decrease with diffusion rate (Fig. S2).

**Figure 2.**
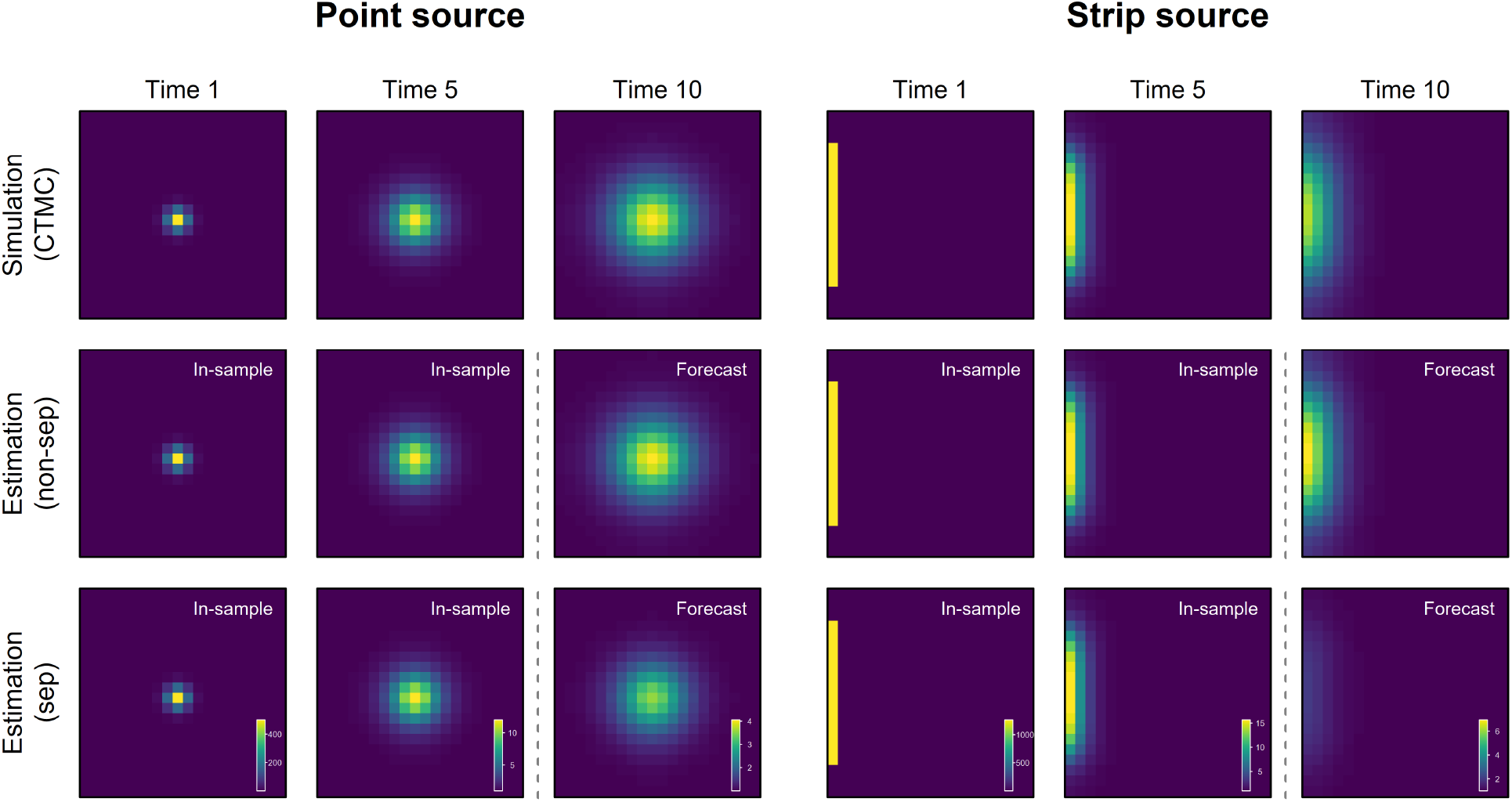
Visualizing CTMC-simulated diffusion dynamics (top row) and corresponding approximations from the non-separable (middle row) and separable (bottom row) DESTMs at selected time steps, shown from left to right (*t* = 1, 5, and 10). Both models were fitted using CTMC-simulated data from *t* = 1–7 (in-sample) and projected to *t* = 8–10 (forecast). A common color scale was used across models within each time step. The weak signal in the separable forecast at *t* = 10 reflects the decay in the spatio-temporal effect over the forecast period.

**Figure 3.**
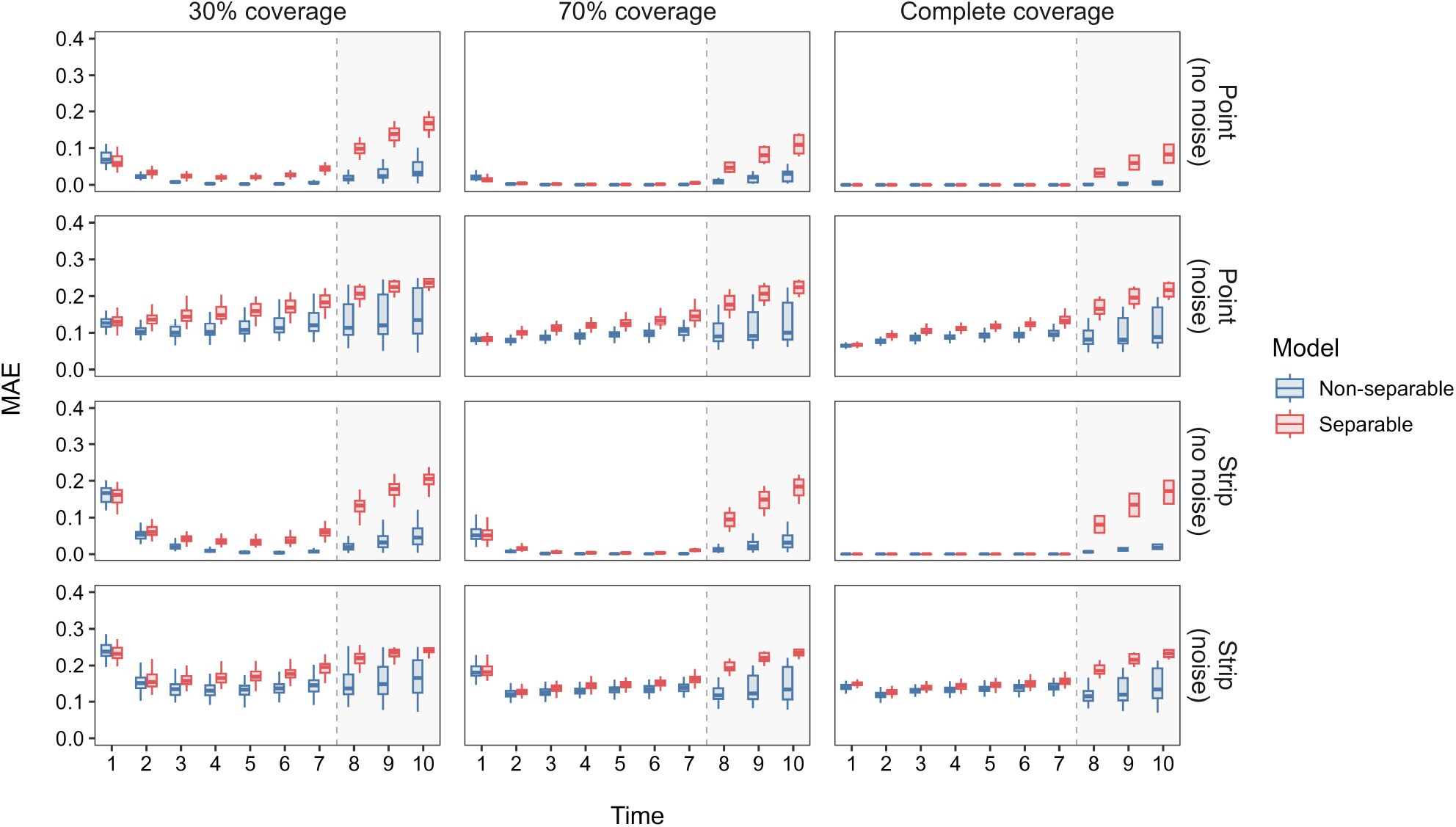
Model performance of non-separable (blue) and separable (red) DESTM fitted to CTMC-simulated diffusion data under four simulation scenarios: point-source with fully observed data (first row), point-source with multinomial sampling (second row), strip-source with fully observed data (third row), and strip-source with multinomial sampling (fourth row). Columns show three observation coverage levels (30%, 70%, and 100%). White regions indicate in-sample estimation (*t* = 1–7), and shaded gray regions indicate forecast periods (*t* = 8–10).

To further evaluate model behavior, we simulated data with and without diffusion using the non-separable and separable DESTMs as operating models (Fig. 4). The non-separable estimation model provided better overall performance, including both in-sample prediction (left panel) and forecasting (right panel), when diffusion was present. In contrast, both models performed similarly for in-sample prediction (left panel) and forecasting (right panel) when diffusion was absent, suggesting that the non-separable estimation model can revert to the separable model while still capturing diffusion when it occurs.

**Figure 4.**
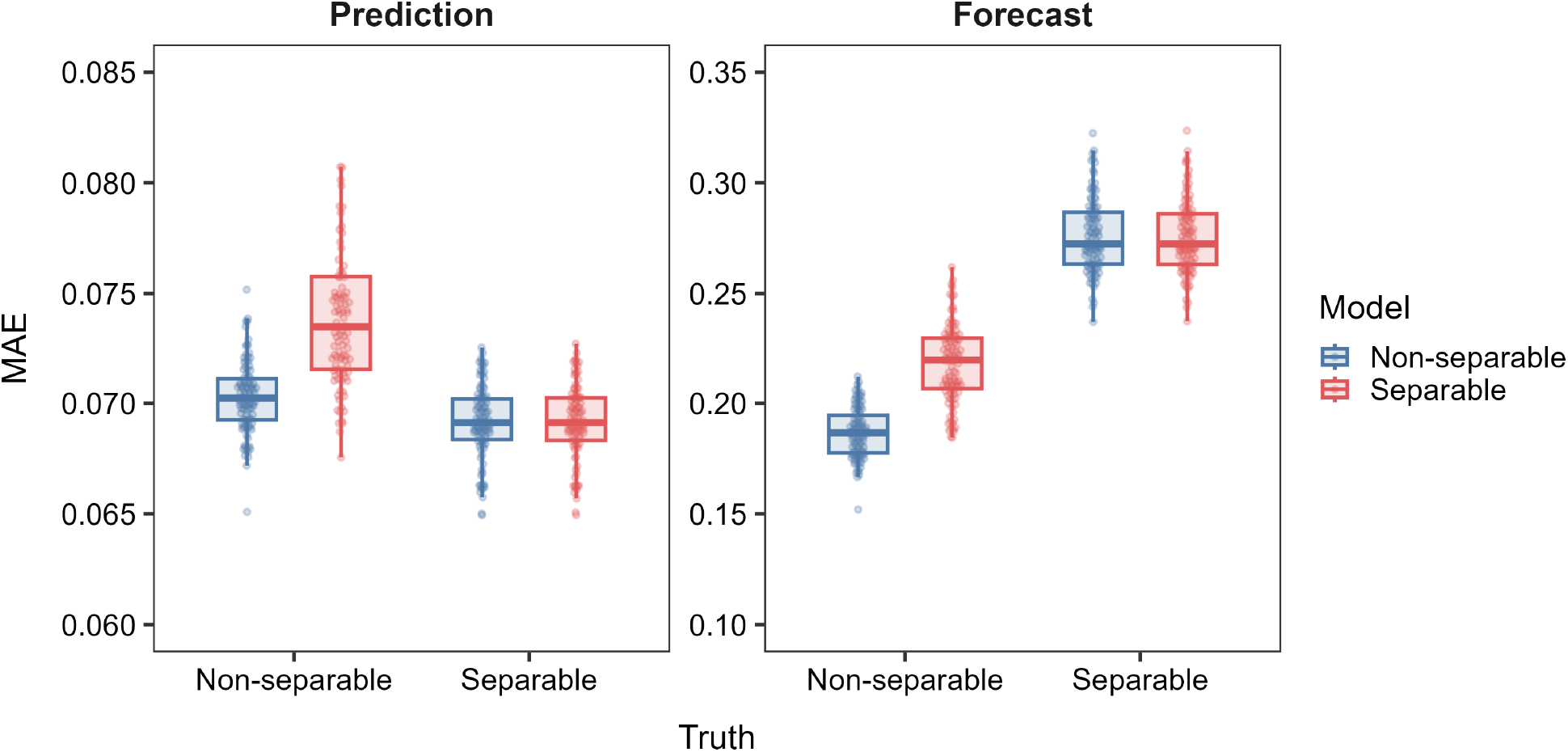
Model performance of non-separable (blue) and separable (red) DESTM under self- and cross-fitting experiments. Datasets were simulated from separable and non-separable operating models (x-axis) using parameter estimates obtained from the CTMC simulations, and fitted using both estimation models. Prediction (left panel) and forecast (right panel) performance are summarized by mean absolute error (MAE).

### 3.2. Japanese distant-water longline fishery

We next applied the DESTM to the historical expansion of the Japanese longline fishery in the Pacific Ocean. The non-separable model was more parsimonious than the separable model (ΔmAIC = 96.3; Table 1) and had higher conditional deviance explained and COR, as well as lower MAE, RMSE, and CRPS. To illustrate the fitted spatio-temporal dynamics, we visualized the estimated fishing effort log(*µ*_*s,t*_), and diffusion process *ϵ*_*s,t*_ for Equation (16) for the mAIC-selected non-separable model for selected years (Fig. 5). Predicted fishing-effort density and estimated diffusion effects for all years are shown in Figures S3 and S4, respectively. The predicted fishing-effort density showed the historical eastward expansion of the Japanese longline fishery from waters east of Japan into the western Pacific, followed by a gradual decline in overall fishing effort during the early 1970s (top row, Fig. 5). The estimated diffusion effect revealed a persistent eastward expansion from the western Pacific toward the central and eastern Pacific throughout the study period (bottom row, Fig. 5).

**Table 1.**
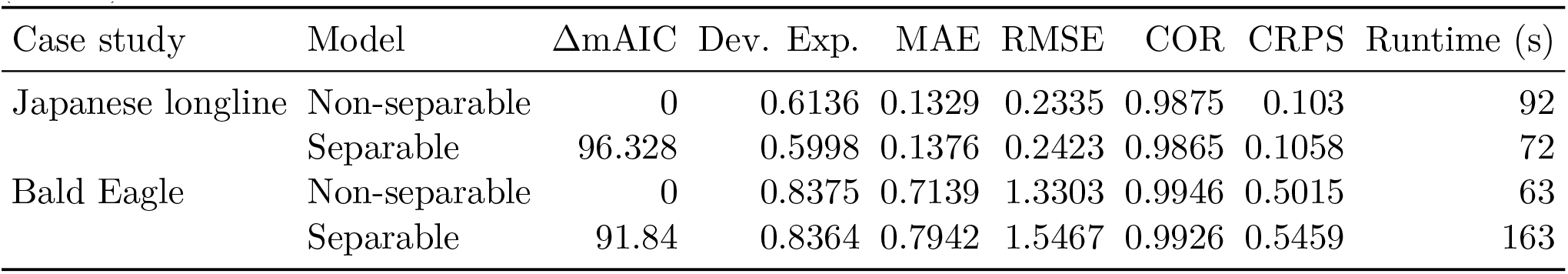
Comparison of non-separable and separable DESTMs fitted to the complete Japanese longline and Bald Eagle datasets. Model performance was evaluated using the difference in marginal Akaike information criterion (ΔmAIC), deviance explained (Dev. Exp.), mean absolute error (MAE), root mean squared error (RMSE), Pearson correlation (COR), continuous ranked probability score (CRPS), and computational runtime in seconds.

| Case study | Model | $\Delta\text{mAIC}$ | Dev. Exp. | MAE | RMSE | COR | CRPS | Runtime (s) |
| --- | --- | --- | --- | --- | --- | --- | --- | --- |
| Japanese longline | Non-separable | 0 | 0.6136 | 0.1329 | 0.2335 | 0.9875 | 0.103 | 92 |
|  | Separable | 96.328 | 0.5998 | 0.1376 | 0.2423 | 0.9865 | 0.1058 | 72 |
| Bald Eagle | Non-separable | 0 | 0.8375 | 0.7139 | 1.3303 | 0.9946 | 0.5015 | 63 |
|  | Separable | 91.84 | 0.8364 | 0.7942 | 1.5467 | 0.9926 | 0.5459 | 163 |

**Figure 5.**
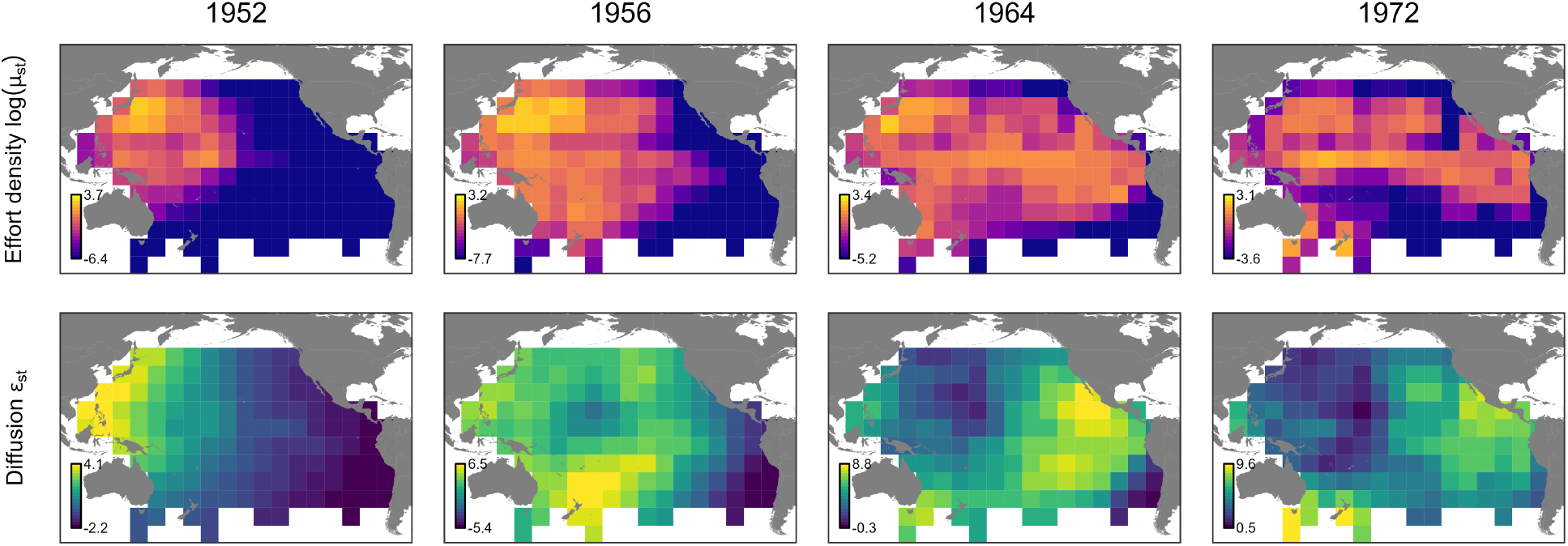
Predicted effort density log(*µ*_*s,t*_) (top row), and estimated diffusion effects *ϵ*_*s,t*_ (bottom row) from Equation 15 for the mAIC-selected non-separable DESTM for the historical Japanese tuna longline fishery in the Pacific Ocean in selected years for 1952, 1956, 1964, and 1972. The diffusion effects show the progressive eastward and southward expansion of Japanese longline fishing effort over time, beyond the time-invariant spatial effect *ω*_*s*_.

To illustrate how the DESTM forecasts spatial expansion in practice, we present one example of a rolling-window forecast using observations from 1952–1957 to predict fishing effort over the following three years (1958, 1959, and 1960; Fig. 6). Observed fishing effort (first row) is directly compared with the corresponding fitted and forecast fishing effort from the non-separable (middle row) and separable (bottom row) models. Both models provided similar in-sample predictions for the final fitted year (1957, first column). However, differences became increasingly apparent during forecasting. The non-separable model prediction maintained the observed eastward expansion and tracked the expansion front (white contour, representing a common effort-density threshold) over the following three years. In contrast, the separable model seemed to underestimate the spatial expansion, particularly in the second- and third-year forecasts. We then summarized forecast performance across all rolling-window analyses to evaluate overall forecasting skill throughout the study period (Fig. 7). Forecast errors generally increased with forecast horizon for both models, as expected. Across all forecast horizons, however, the non-separable model consistently achieved lower MAE, RMSE, and CRPS and higher correlation than the separable model, with the advantage becoming more apparent at longer forecast horizons.

**Figure 6.**
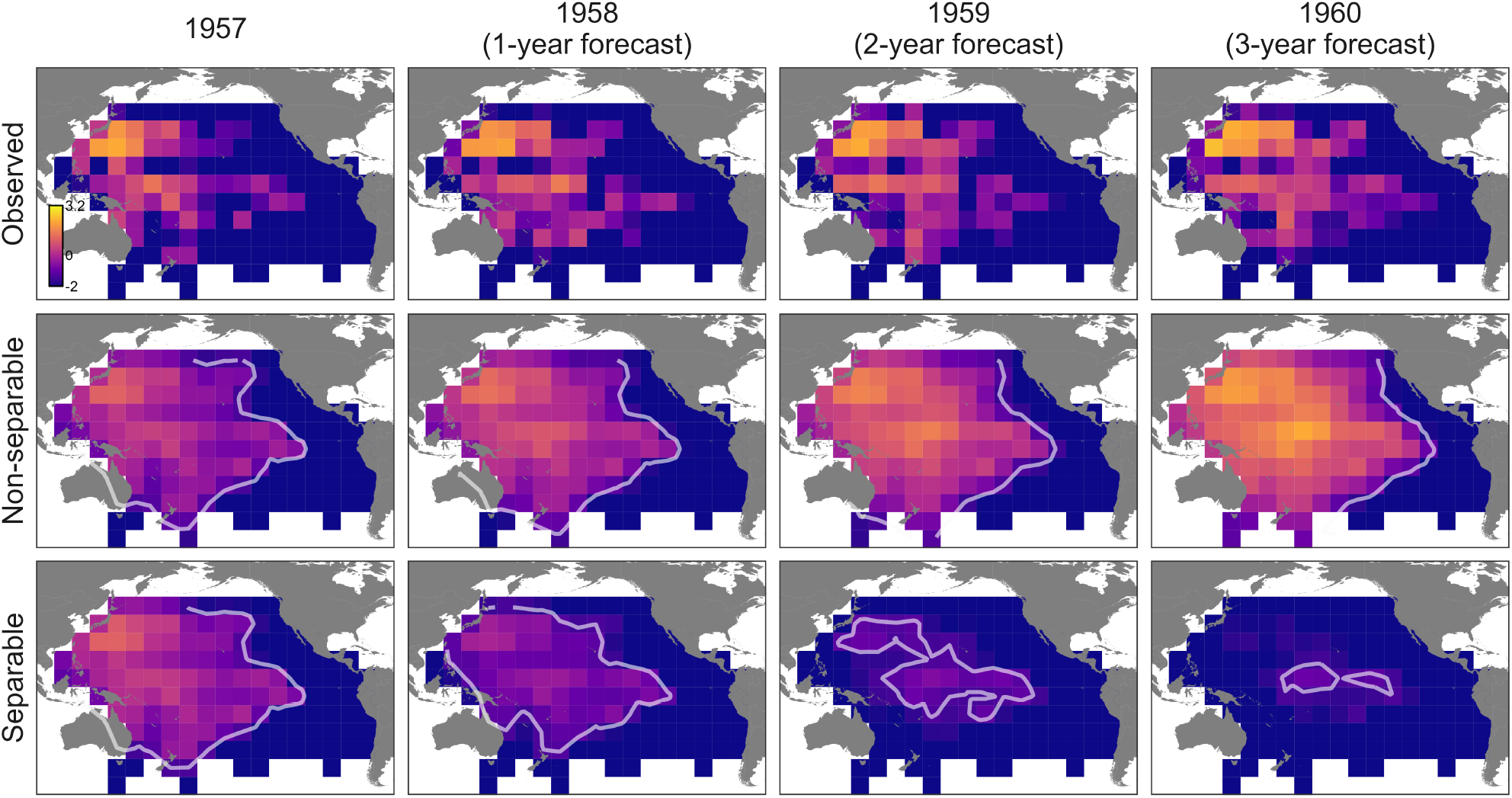
An example rolling-window forecast for the Japanese tuna longline fishery, fitted using observations from 1952–1957 and showing fitted fishing effort for 1957 (final fitting year, first column). The fitted models were then used to forecast effort expansion for 1958 (first-year forecast, second column), 1959 (second-year forecast, third column), and 1960 (third-year forecast, fourth column). Observed effort density (top row) is compared with predictions from the non-separable DESTM (middle row) and separable DESTM (bottom row). White contours indicate the estimated expansion front, defined as the boundary of fishing activity corresponding to a fixed effort-density threshold of 0.25.

**Figure 7.**
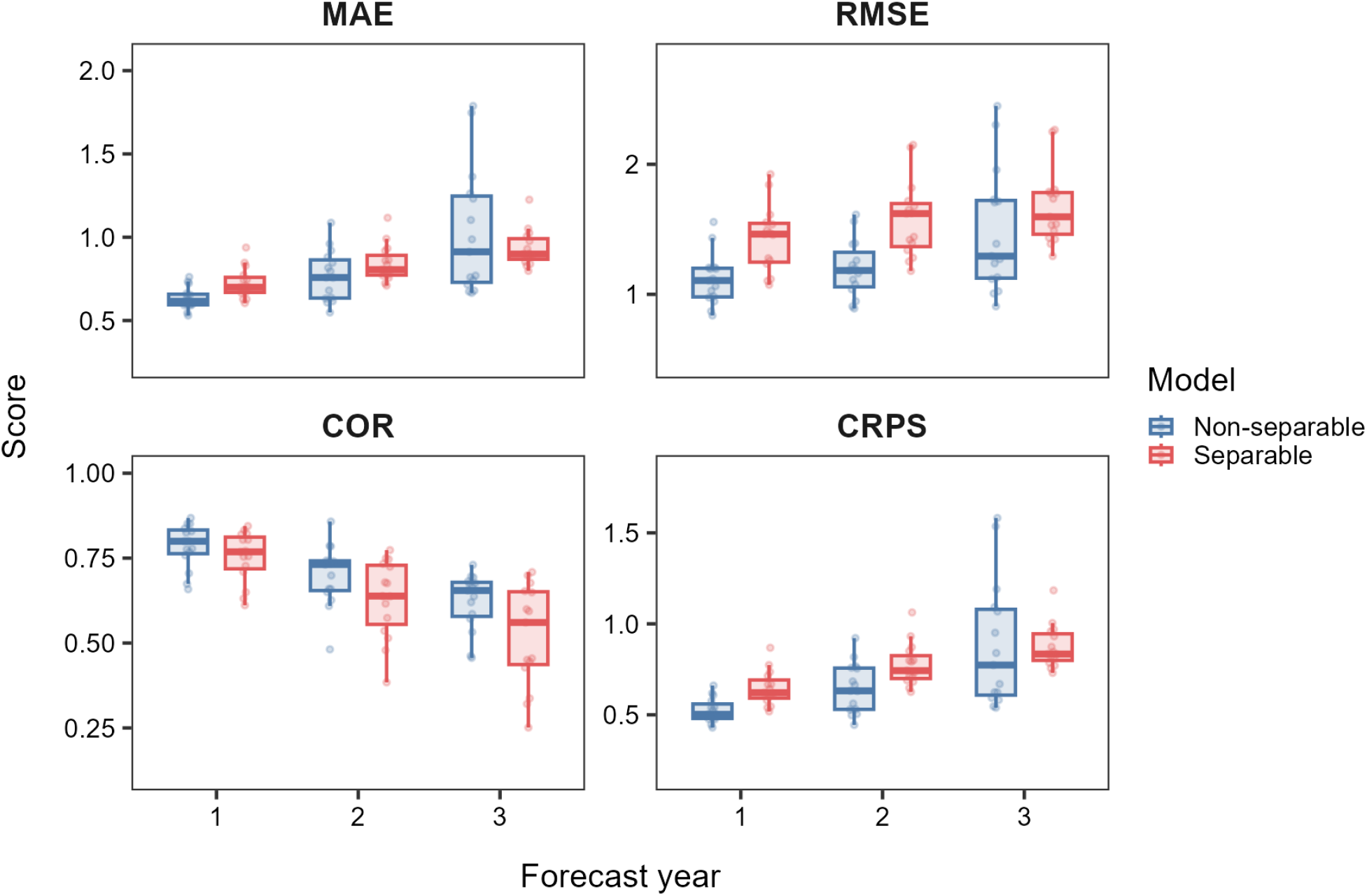
Forecast skill of non-separable (blue) and separable DESTMs (red) evaluated using leave-future-out cross-validation with an rolling window for the Japanese tuna longline fishery. Models were fitted using observations up to year *t* and used to forecast fishing effort for years *t* + 1, *t* + 2, and *t* + 3, with forecasts repeated through the final evaluation period (1972–1974). Forecast performance was evaluated using mean absolute error (MAE), root mean squared error (RMSE), correlation (COR), and continuous ranked probability score (CRPS).

### 3.3. Bald Eagle population recovery

Finally, we applied the DESTM to the recovery and geographic expansion of Bald Eagle populations across North America. Predicted relative density and estimated diffusion effects from the non-separable model throughout the study period are shown in Figures S5 and S6, respectively. Model comparison showed that the non-separable model was more parsimonious than the separable model based on marginal AIC (ΔmAIC=91.84; Table 1). The non-separable model also had lower MAE, RMSE, and CRPS, although differences in deviance explained and COR were small. However, this improvement in full-data fit did not translate into a clear forecast advantage, and the two models showed similar performance in the leave-future-out forecasts (Fig. 8). Forecast performance declined with forecast horizon for both models, with increasing RMSE, MAE, and CRPS and decreasing COR. The non-separable model tended to have lower RMSE and CRPS and slightly higher COR, particularly at longer forecast horizons, while neither model consistently had lower MAE (Fig. 8, top panels). Both models predicted higher probabilities for observed presences than for observed absences, but the non-separable model tended to predict higher probabilities for both groups (Fig. 8, bottom panels).

**Figure 8.**
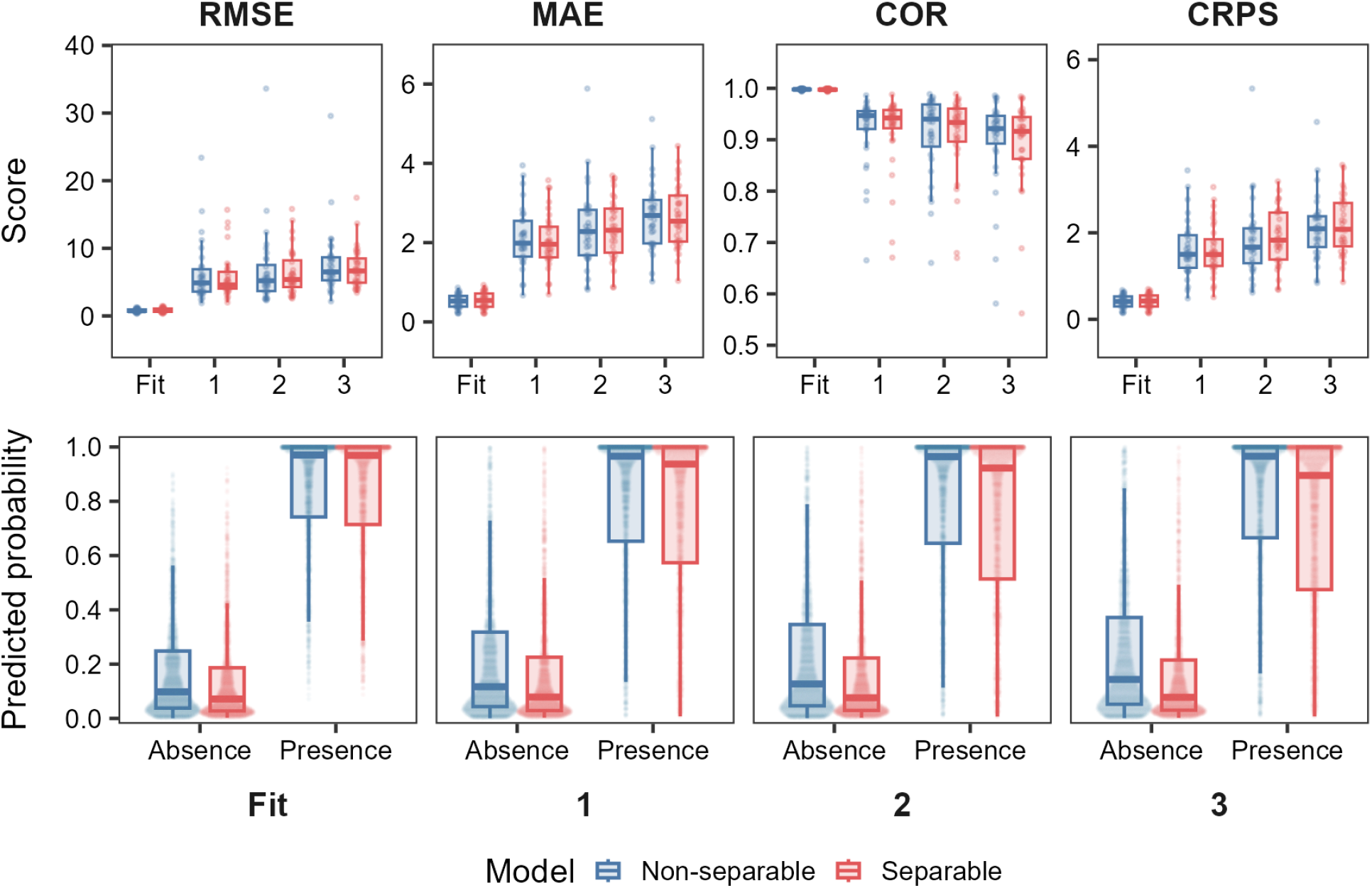
Leave-future-out cross-validation results for the Bald Eagle case study comparing the non-separable (blue) and separable (red) DESTMs. Models were fitted using observations up to year *t* and evaluated for the final fitted year *t* and forecasts for years *t* + 1, *t* + 2, and *t* + 3. Forecast performance was summarized using root mean squared error (RMSE), mean absolute error (MAE), Pearson correlation (COR), and continuous ranked probability score (CRPS) (top panels). Predicted probabilities for observed absences and presences are shown for each evaluation horizon (bottom panels).

## 4. Discussion

In this paper, we introduced diffusion into conventional spatio-temporal models to enable latent diffusion processes to be estimated directly from ecological observations. The DESTM represents diffusion through a joint spatio-temporal precision structure in which the latent state at one time step propagates to neighboring locations at the next. In simulations, the model closely approximated mechanistic diffusion dynamics and produced lower estimation and forecast errors than the separable model when diffusion was present. When diffusion was absent, it reverted to a separable form, showing that it can accommodate conditions ranging from no diffusion to pronounced diffusion. We then applied the DESTM to two examples of spatial expansion: the historical spread of Japanese longline fishing effort across the Pacific and Bald Eagle population recovery across North America. The non-separable model was more parsimonious in both applications, but improvements in forecast performance were more apparent for the longline fishery. In the longline application, the inferred latent process also captured the progressive expansion of fishing effort. The results show that the DESTM can estimate diffusion directly and use the same latent process to predict and forecast spatial expansion, although its forecasting advantage over the separable model varied between applications.

We highlight an important distinction between conventional spatio-temporal statistical models and DESTMs. In conventional STMs, latent random fields are used to account for residual spatial and temporal dependence that is not explained by the fixed effects (Diggle et al., 1998; Latimer et al., 2006). In particular, STMs commonly model spatio-temporal effects by linking spatial fields across time through autoregressive processes (as in the separable model considered here) or random walks, allowing the preceding latent field to be carried forward at the same locations (Thorson, 2019). These flexible dependence structures can improve interpolation and forecasting at unsampled locations or times, but the dependence captured by the estimated random fields cannot always be attributed to a specific ecological process (Cressie and Wikle, 2011; Wikle and Hooten, 2010). This limitation may also affect forecasting because such STMs can project the persistence or decay of existing spatial effects but do not explicitly describe how the latent state propagates among neighbouring locations (Fig. 2 and 3). By contrast, the DESTM links the latent state at each location to neighbouring locations at the preceding time within the joint precision structure. This constrains the latent dependence to follow local, temporally ordered propagation, allowing an existing spatial field to spread rather than only persist or decay in place. The DESTM therefore retains the statistical structure of a conventional STM while allowing its latent field to represent diffusive dynamics.

Diffusion has also recently been used to model non-local environmental effects, allowing covariate effects to extend across nearby locations and previous times (Lindmark et al., 2026; Thorson et al., 2026). In the DESTM, diffusion instead directly accounts for the propagation of the latent state among locations through time. Diffusive dynamics can also be modeled more explicitly using process-based approaches, often through partial differential equations (PDEs) that describe how an ecological state changes across space and time (Hefley et al., 2017a; Wikle and Hooten, 2010). Process models can then be linked to probabilistic observation models, allowing ecological process parameters to be estimated while accounting for observation uncertainty within a mechanistic–statistical model (Louvrier et al., 2020; Soubeyrand and Roques, 2014; Williams et al., 2017). However, this explicit process specification also requires defining the model structure, parameterization, initial conditions, and boundary conditions, which can become challenging for high-dimensional or finely resolved spatial domains. (Hefley et al., 2017a; Soubeyrand and Roques, 2014; Wikle and Hooten, 2010). Diffusion equations can also be used within the latent statistical process; for example, advection–diffusion SPDE models define Gaussian processes through stochastic process equations, allowing diffusion and directional transport to shape latent spatio-temporal dependence (Berild and Fuglstad, 2024; Clarotto et al., 2024; Sigrist et al., 2015). In the DESTM, diffusion is specified in a simpler, graph-based form by embedding a lagged diffusion operator in the precision matrix of the latent GMRF. Initial conditions are estimated as latent fields, while the spatial graph and zero-row-sum diffusion operator imply a no-flux boundary, i.e., restricting diffusion to the modeled domain and conserving the total within it. The resulting latent dynamics have a clearer process interpretation than those of a conventional latent random field while requiring less detailed process specification than a fully mechanistic model.

We note that the DESTM assumes a stationary diffusion process throughout the fitted time series, with both local temporal dependence and spatial diffusion remaining constant through time. In addition, the DESTM assumes symmetric diffusion and does not explicitly include local reaction dynamics. When expansion dynamics vary through time, for example because dispersal or demographic processes change during range expansion (Chuang and Peterson, 2016; Kubisch et al., 2014; Miller et al., 2020), these changes may instead be absorbed by time-specific latent variation. These assumptions could be relaxed to allow the latent spatio-temporal process to accommodate additional ecological dynamics. For example, stationary path parameters could be replaced with time-varying parameters to accommodate temporal nonstationarity using period-specific parameters or autoregressive processes (Rollinson et al., 2021; Thorson and Kristensen, 2026). Likewise, symmetric diffusion could be relaxed by extending the spatial operator to include asymmetric weights representing passive advection or taxis along habitat gradients (Thorson et al., 2021). Explicit local reaction dynamics could also be incorporated by extending the temporal correlation beyond linear carry-over to account for density-dependent growth, natural mortality, or harvesting (Abboud et al., 2019; Thorson et al., 2017). Beyond these methodological extensions, integrating the DESTM into widely used spatio-temporal modelling software, such as sdmTMB (Anderson et al., 2025) or tinyVAST (Thorson et al., 2025), would provide a practical platform for these developments and make the framework accessible to a broader range of ecological applications.

## Acknowledgments

We thank Sean C. Anderson and Federico Maioli for helpful comments on an earlier draft.

## Data Availability

The Japanese longline fleet data used in this study are publicly available from the Western and Central Pacific Fisheries Commission (WCPFC) Public Domain Aggregated Catch/Effort Data (https://www.wcpfc.int/public-domain-aggregated-catcheffort-data; downloaded February 2025) and the Inter-American Tropical Tuna Commission (IATTC) Public Domain Data for download (https://www.iattc.org/en-US/Data/Public-domain; downloaded February 2025). The North American Breeding Bird Survey (BBS) data are publicly available from the U.S. Geological Survey BBS data (https://www.pwrc.usgs.gov/bbs/RawData/; downloaded July 2026). All code used for will be made publicly available on GitHub unpon publication at https://github.com/James-Thorson-NOAA/diffusion-enhanced.

