## Supplementary Figures for "Diffusion-enhanced spatio-temporal models for estimating spatial expansion"

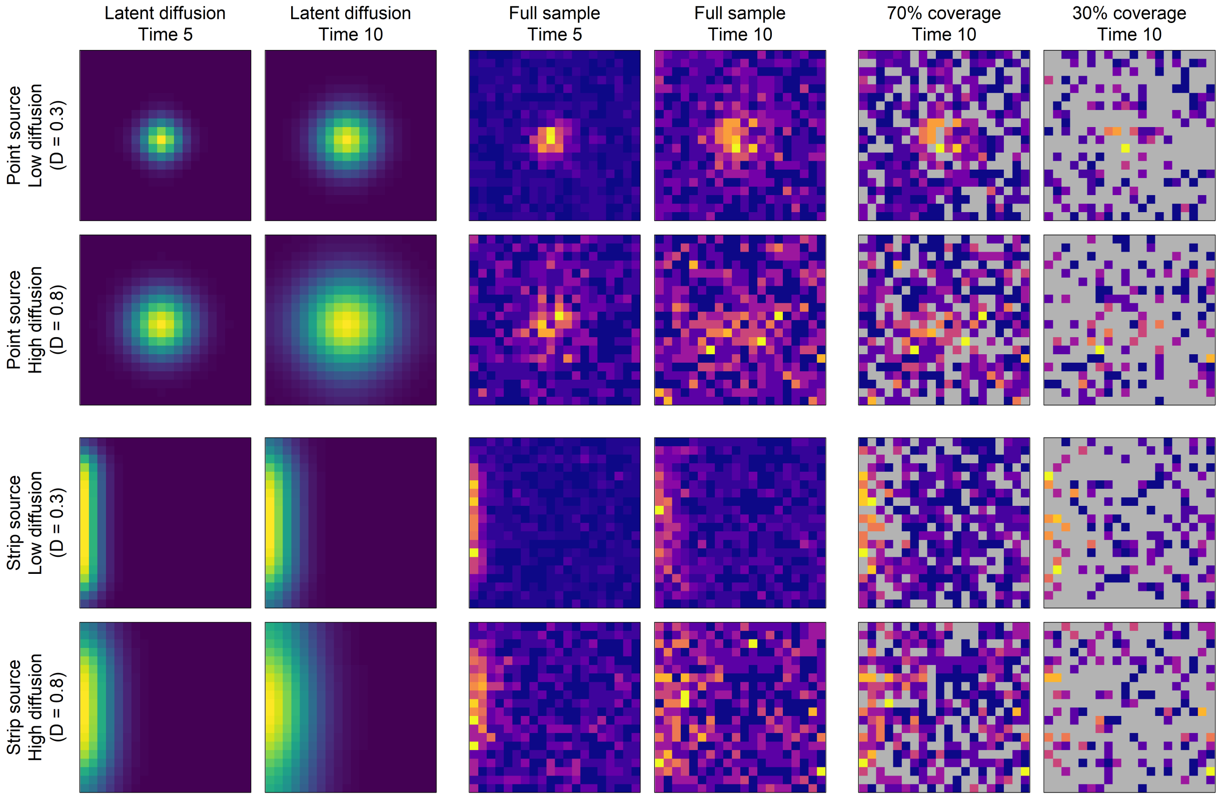


Figure S1: CTMC diffusion scenarios used in the simulation experiment. We show point-source simulations (top panels) and strip-source simulations (bottom panels), each under low ($\delta$=0.3) and high ($\delta$=0.8) diffusion rates. Latent diffusion is shown at t=5 and t=10 (first and second columns), followed by full samples at the same time steps (third and fourth columns) and samples with 70% and 30% observation coverage at t=10 (fifth and sixth columns). Full samples included process error and were generated using a multinomial distribution. For each scenario, the samples with 70% and 30% observation coverage were all based on the same full sample, with gray cells indicating locations without observations. Color scales were mapped separately to the value range in each panel to make the spatial patterns visible.

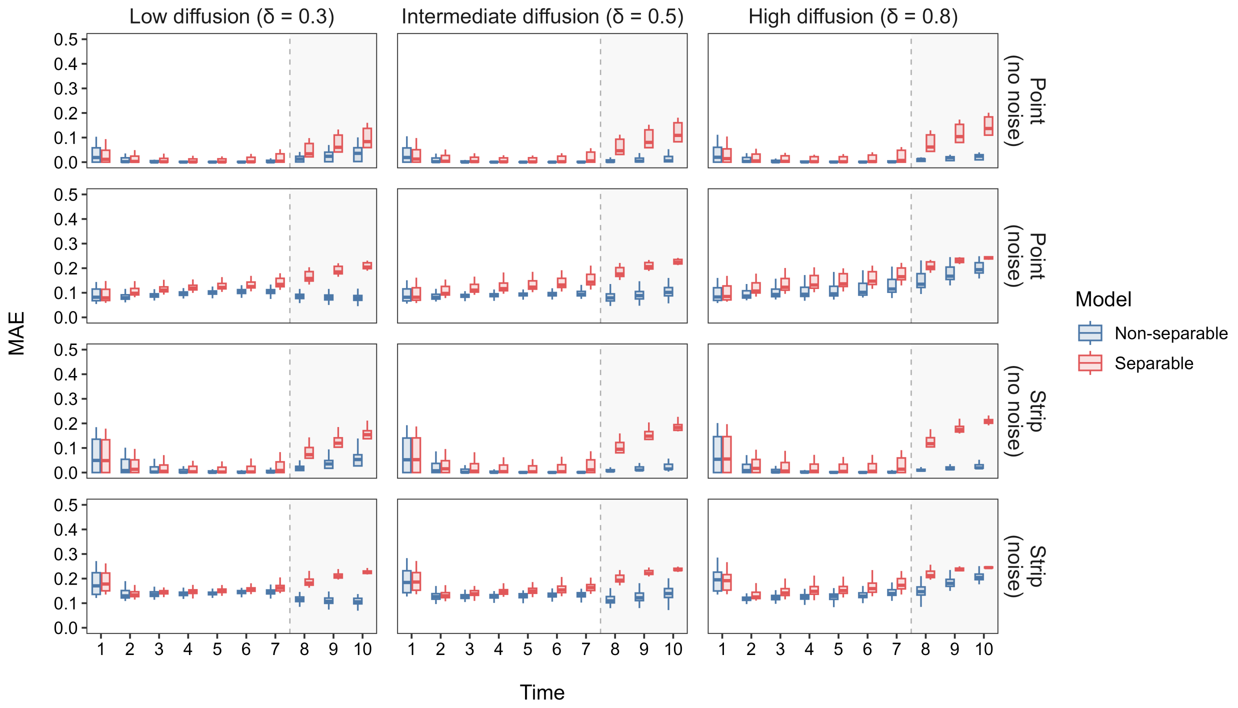


Figure S2: Model performance of the non-separable (blue) and separable (red) DESTMs fitted to CTMC-simulated diffusion data under four simulation scenarios: point-source without process error (first row), point-source with multinomial sampling (second row), strip-source without process error (third row), and strip-source with multinomial sampling (fourth row) for low ($\delta = 0.3$; first column), intermediate ($\delta = 0.5$; second column), and high ($\delta = 0.8$; third column) diffusion rates. Performance was evaluated using mean absolute error (MAE), with each boxplot combining 30 simulation replicates from each of the three observation coverage levels (30%, 70%, and 100%). White regions indicate in-sample estimation (t = 1–7), and shaded gray regions indicate forecast periods (t = 8–10).

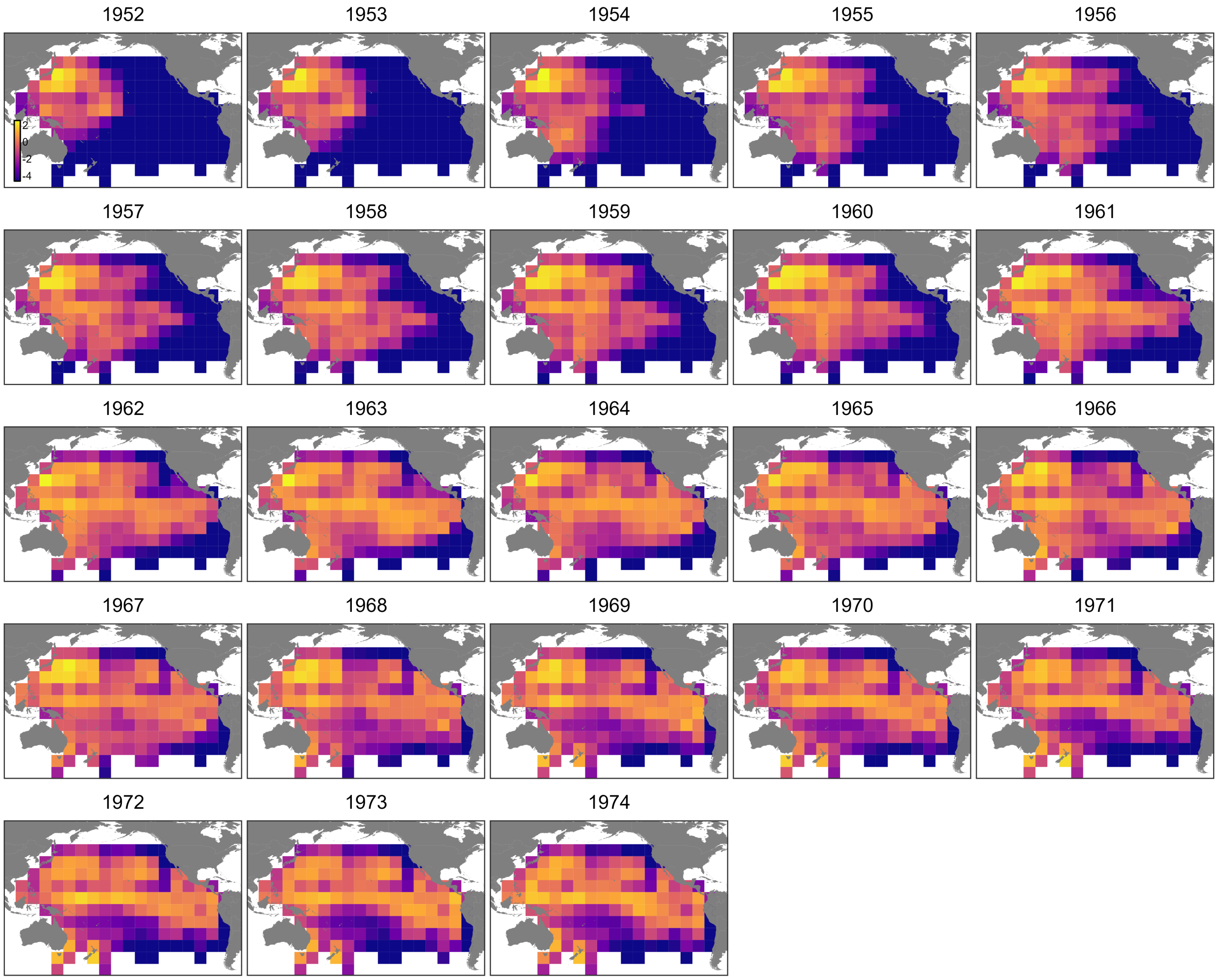


Figure S3: Distributions of predicted fishing-effort density on the log scale $\log(\mu_{s,t})$ in Equation (16) for the Japanese tuna longline fishery using the AIC-selected non-separable DESTM in the Pacific Ocean from 1952 to 1974.

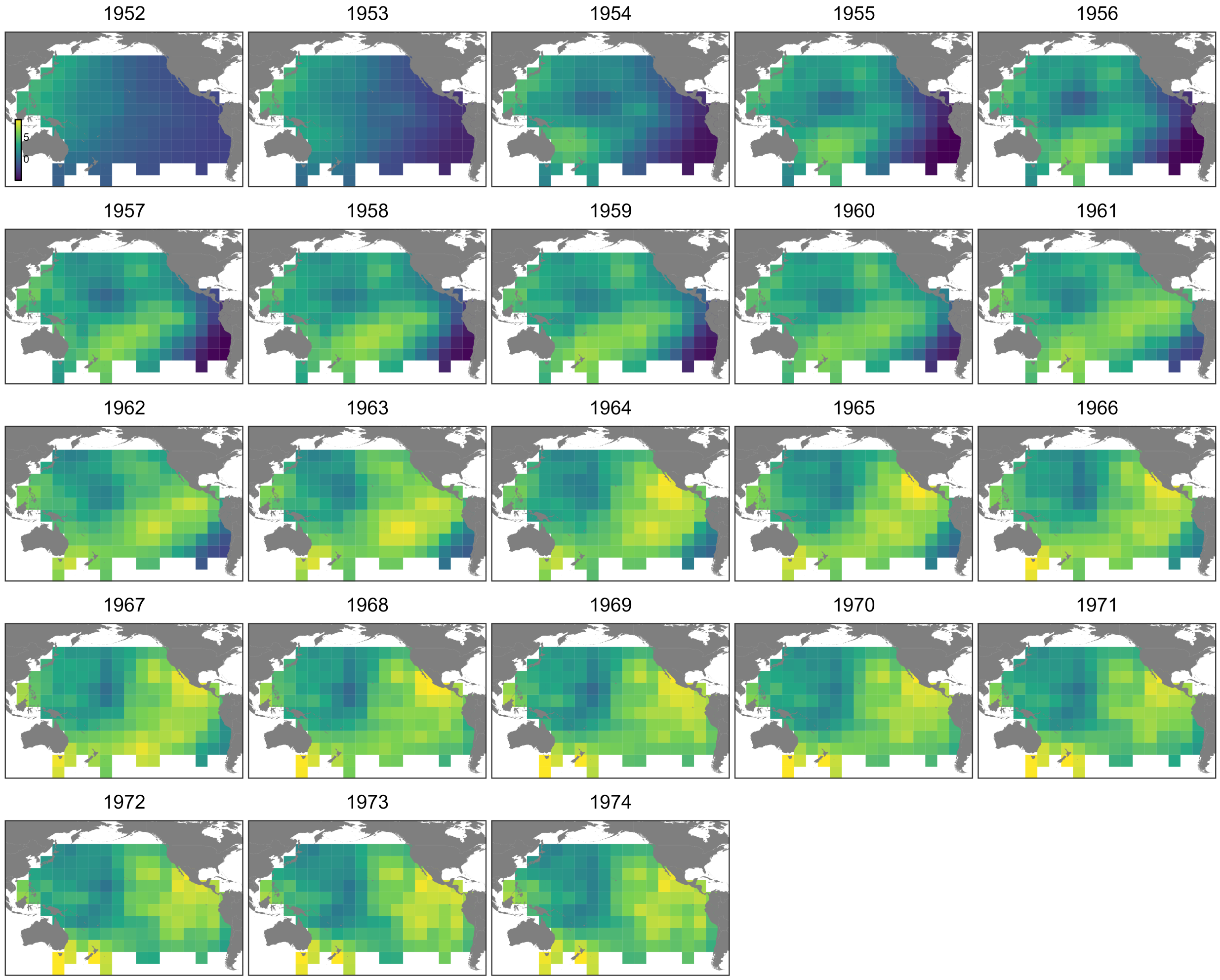


Figure S4: Estimated diffusion effects $\epsilon_{s,t}$ in Equation (16) for the Japanese tuna longline fishery using the AIC-selected non-separable DESTM across the Pacific Ocean from 1952 to 1974.

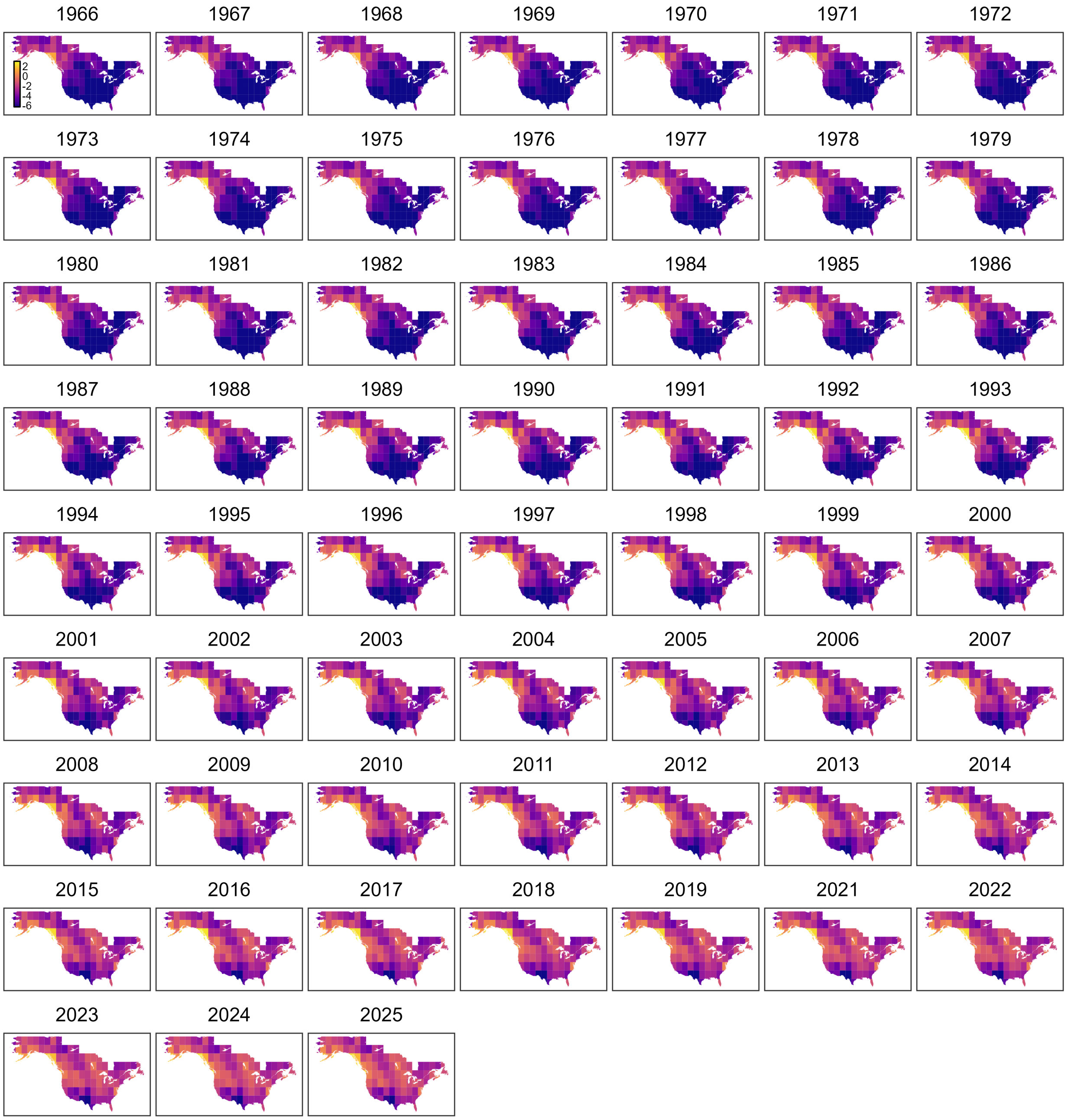


Figure S5: Distributions of predicted relative density on the log scale [$\log(\mu_{s,t})$, in Equation (21)] for Bald Eagle from the AIC-selected non-separable DESTM in North America from 1966 to 2025.

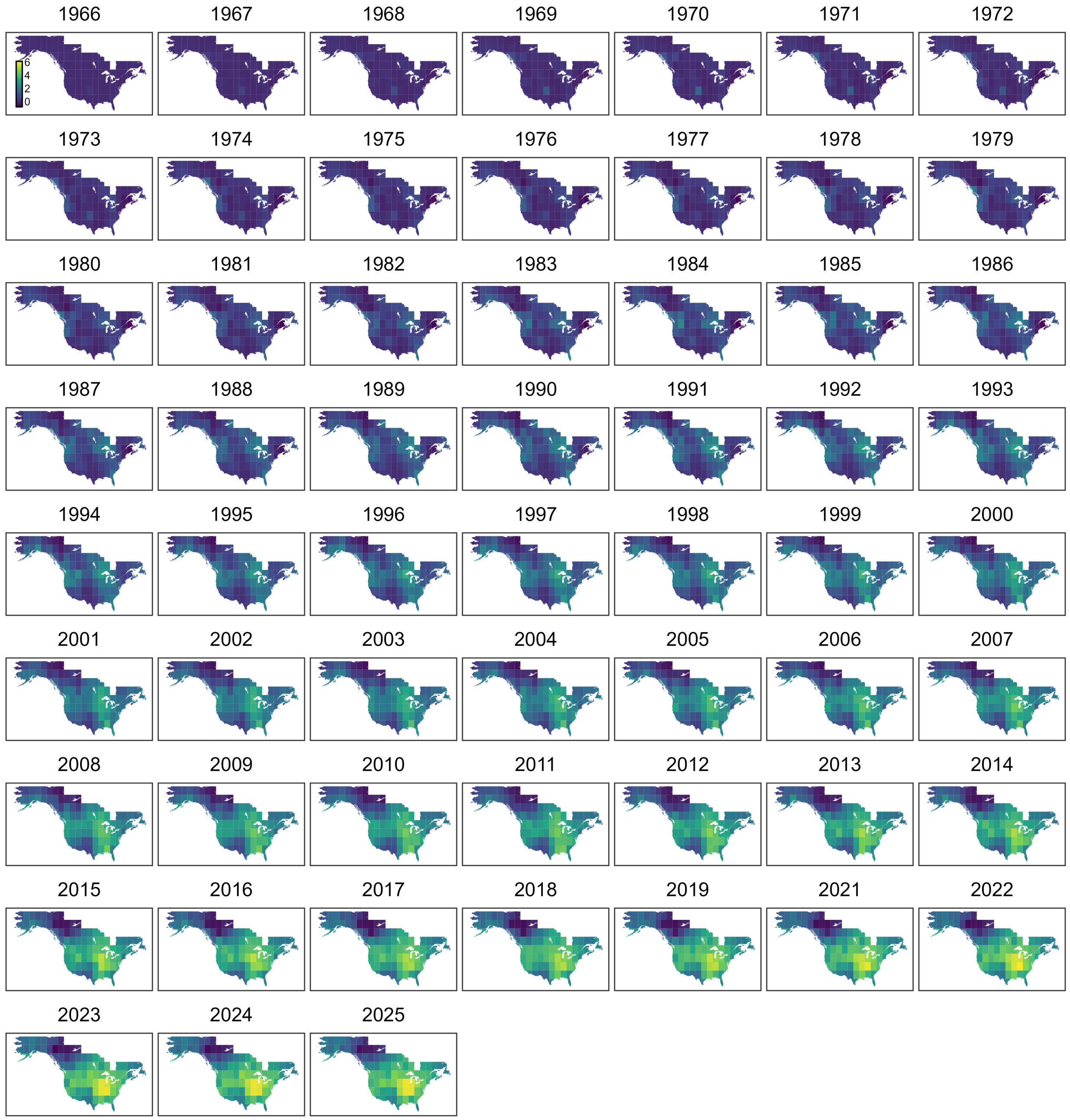


Figure S6: Estimated diffusion effects [$\epsilon_{s,t}$, in Equation (21)] for Bald Eagles using the AIC-selected non-separable DESTM in North America from 1966 to 2025.
